# Combining local knowledge from oral histories and participatory mapping with veteran fishers to identify long-term environmental change in a nationally significant river

**DOI:** 10.64898/2025.12.26.695674

**Authors:** Shane Orchard, Oakley Campbell

## Abstract

The detection and monitoring of environmental change is vital for sustainable river management, but historical data are often limited. This study leveraged volunteered geographic information and local ecological knowledge from experienced recreational anglers to identify historical baselines and long-term change in the Rakaia River, a protected catchment under a National Water Conservation Order (WCO) in Aotearoa New Zealand. Oral histories spanning seven decades were recorded using semi-structured interviews with 30 veteran fishers representing 1510 years of combined catchment-specific experience. Inductive thematic analysis and participatory mapping were used to identify physical environment changes and socio-cultural effects associated with recent declines in freshwater fish populations. River environment changes include the loss of natural features such as springs and pools, declines in characteristic native species and habitats, shrinking river mouth la-goon and river plume extents, and interactions with public access. Socio-cultural effects include reduced participation in fishing, community despondence, emotional distress and demographic shifts in fishing hut settlements. Over 20 reported changes involve adverse effects on values that are specifically protected under the WCO, indicative of a policy failure. Declines in multiple indicators of environment health suggest an implementation gap that requires greater attention to environmental monitoring and outcome evaluation yet is hampered by uncertainties around institutional responsibilities. Oral history approaches can help to establish historical baselines for gauging long-term change and enabling adaptive management in this and other data-poor situations.

## 1. Introduction

The measurement of environmental change has a crucial function in sustainable resource management through its role in the feedback loop between people and resources (Lindenmayer & Likens, 2010). It enables practices such as adaptive management and facilitates forward-looking practices such as risk and vulnerability assessment (Folke et al., 2005; Turner et al., 2003). Knowledge of what once existed is both an indicator of societal understanding and an important reference point for goal setting and future evaluations (Dayton et al., 1998). Maintaining this knowledge also helps to prevent the shifting baseline syndrome, where institutional or societal memory is lost leading to downwards adjustments in environmental outcomes and expectations (Papworth et al., 2009; Pauly, 1995).

Participatory science offers a solution where members of the public contribute to collective knowledge through the sharing of observations (Dickinson et al., 2010; Shirk et al., 2012). Over history, the act of making observations has long supported environmental awareness, understanding and connection (Jopling et al., 2024; Tolbert et al., 2024; Toomey & Domroese, 2013). It can also assist communities to navigate the complexities of environmental change by supporting a shared understanding and interest in changing conditions. In this context, the observations of community members who have lived through significant historical events represents a unique source of knowledge that has great potential to assist the understanding of former conditions (Miller-Rushing et al., 2012; Silvertown, 2009). In this study we explored this potential by developing a participatory approach with veteran members of fishing communities to assess social-ecological change in the Rakaia River catchment in Aotearoa New Zealand (NZ).

The Rakaia is one of several large, braided river catchments that are defining features of the Waitaha / Canterbury Region and have been heavily modified by water abstraction (Figure 1). However, it is also subject to a National Water Conservation Order (WCO), which is a statutory instrument that provides a high level of protection for the outstanding amenity or intrinsic values of water bodies (Box 1) (New Zealand Government, 1988). The status conferred has similarities with statutory recognition mechanisms in the USA under the National Wild and Scenic Rivers System (Public Law 90-542; Congress 1968). As with many NZ rivers, exotic sportfish species have been introduced to create recreational fisheries that are specifically protected under the Rakaia WCO (New Zealand Government, 1988). Despite this, there has been a well-documented decline in the Chinook salmon (*Oncorhynchus tshawytscha*) fishery over the period in which the WCO has been in force (Appendix 1, Figure S1) (Dellaway, 2025). Furthermore, a marked decline in the formerly abundant keystone species, Stockell’s smelt (*Stokellia anisodon*), was recently reported in the lower river along with reduced catches of sea run brown trout (Eldon & Greager, 1983; Hickford, 2022; Jellyman & Mayall-Nahi, 2022; McDowall, 1993).

**Figure 1.**
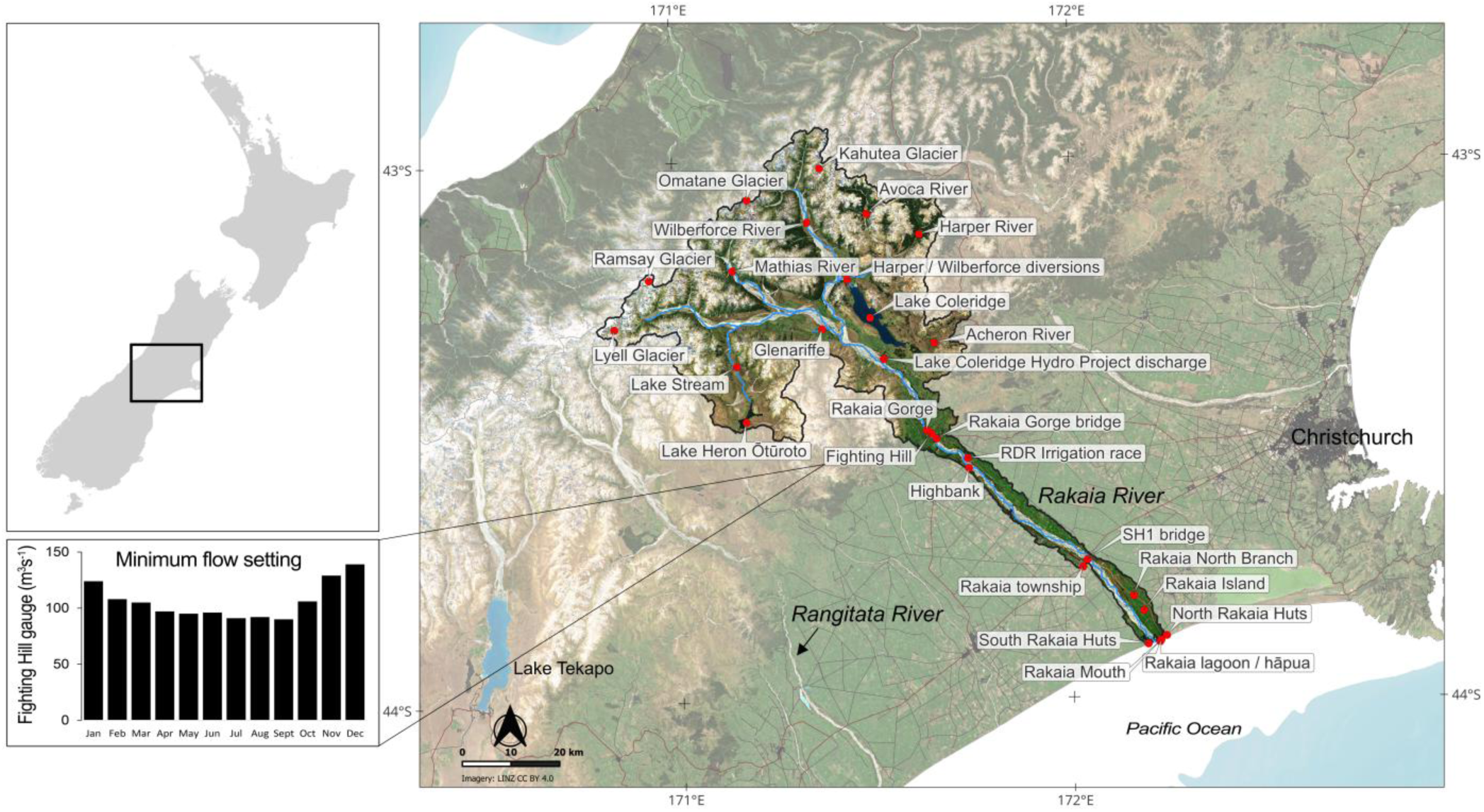
Key locations and geographical context of the Rakaia catchment in the Canterbury Region of Aotearoa New Zealand showing the minimum flow settings required under the National Water Conservation (Rakaia River) Order 1988 as measured at Fighting Hill (New Zealand Government, 1988).

Against this backdrop of decline in the state of fish populations, the primary research questions for this study involved: 1) identifying other aspects of river environment change over the same period, and 2) obtaining an understanding of the social-ecological impacts of these changes. Our specific objectives included understanding the local knowledge of longstanding members of the freshwater fishing community and evaluating the utility of this knowledge for gauging long-term environmental change. In the remainder of the paper, we first describe the social-ecological context of the Rakaia River catchment and the data collection methodology. We then summarise the information shared by the fishing community and evaluate key environment trends and their policy implications under the WCO. We conclude by highlighting the transferability of this approach to similar contexts where baseline information may be widely dispersed despite being essential for environmental impact assessments and the monitoring of long-term change.

#### Box 1 Water Conservation Orders

National Water Conservation Orders (WCOs) are a statutory instrument that is designed to recognise and protect the outstanding values of water bodies in Aotearoa New Zealand. They may be applied to achieve any of the following:

- the preservation as far as possible of the water body’s natural state
- the protection of characteristics which the water body has or contributes to:

o as a habitat for terrestrial or aquatic organisms
o as a fishery
o for its wild, scenic, or other natural characteristics:
o for scientific and ecological values:
o for recreational, historical, spiritual, or cultural purposes:
- the protection of characteristics which any water body has or contributes to, and which are considered to be of outstanding significance in accordance with tikanga Māori.

## 2. Methodology

### 2.1 Study catchment

#### Physical character

The Rakaia is one of the largest braided rivers in NZ, arising in the Southern Alps and flowing in a generally eastward direction to enter the Pacific Ocean in the South Canterbury Bight on the east coast of the South Island. Throughout much of the catchment the river is characterised by multiple shifting channel positions in a wide gravel riverbed (Bowden et al., 1983). The upper catchment is mountainous and includes two major alpine tributary catchments (Mathias and Wilberforce rivers) in addition to the headwater of the Rakaia mainstem (Figure 1). These alpine catchments are glaciated and develop a considerable seasonal snowpack. The river has a mean flow of 206 cubic metres per second (cumecs) as measured at the Fighting Hill gauge in the Rakaia Gorge, a prominent natural feature separating the upper catchment from the lower plains (Table 1). However, combinations of rainfall and seasonal snow melt have a considerable influence on the magnitude and timing of flows. In addition, there are considerable surface water losses to groundwater, particularly across the lower plains downstream of Rakaia Gorge and these losses create uncertainties for river flow and water abstraction calculations (Graham & Clark, 2024; Terrink, 2021). The river discharges to the Pacific Ocean where the river mouth is characterised by an elongated perched lagoon or hāpua as with many rivers on NZ’s mixed-sand gravel coastlines (Kirk, 1991; Measures et al., 2020). These non-tidal lagoons are formed by highly dynamic barrier bars with shifting river mouth positions that are continually re-shaped by large swell and flood events.

**Table 1.**
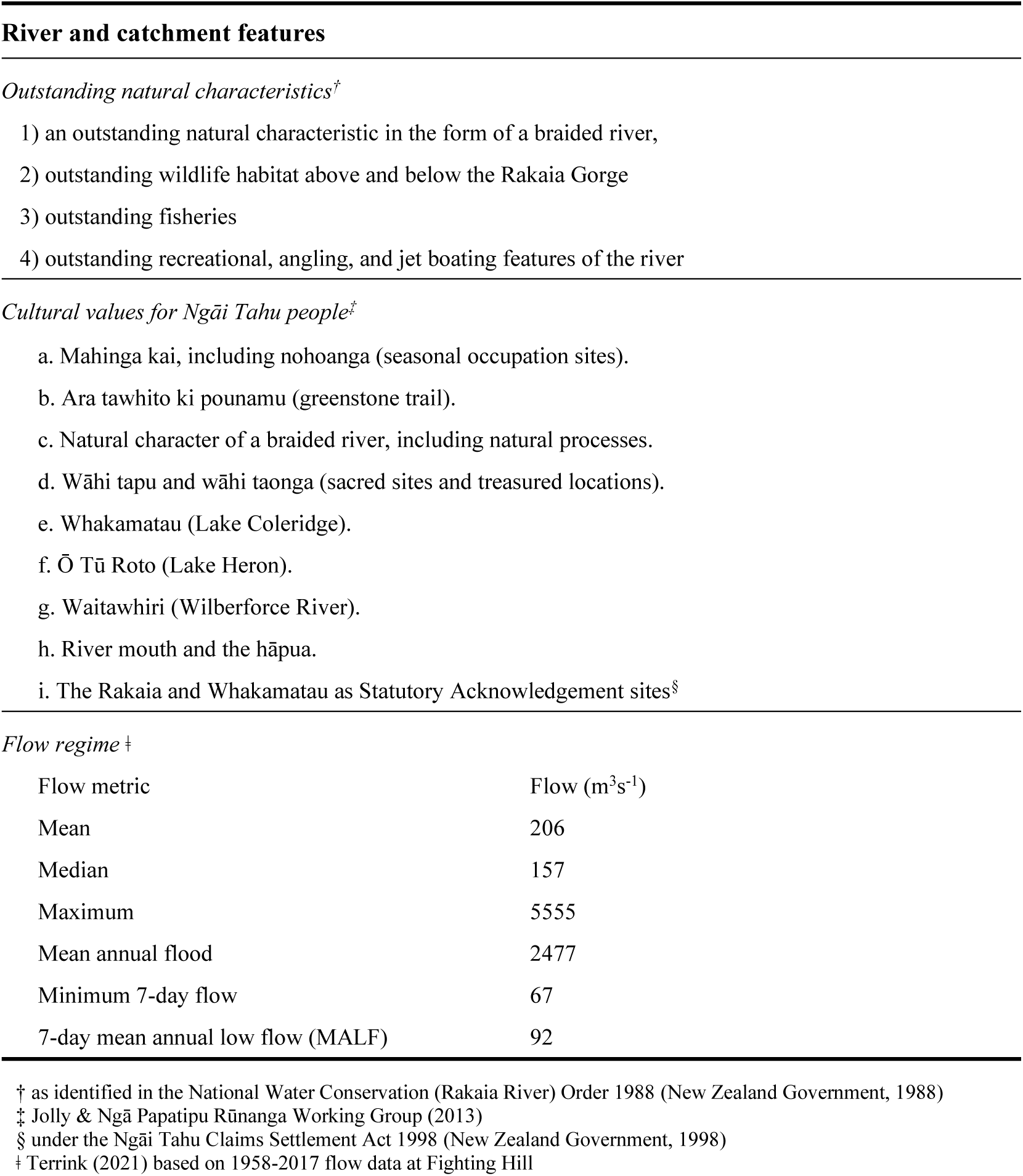
Characteristics of the Rakaia River catchment.

#### Social-ecological context

The Rakaia has significant cultural value for indigenous Ngāi Tahu people (Jolly & Ngā Papatipu Rūnanga Working Group, 2013). In addition to supporting mahinga kai (food gathering) and kaitiakitanga (stewardship) practices of the takiwā (traditional territories), the Rakaia forms the eastern portion of a traditional trail to the West Coast that was an important trade route for pounamu (greenstone) and other resources (Beattie, 1994). The river and associated lake, lagoon and wetland environments provide habitat for native fish and water bird populations and include some of the largest intact wetlands in the Canterbury region (Figure 1). In addition to the introduced sportfish fisheries a wide range of native fish have been recorded in the catchment including several migratory fish that use lagoon and river mouth environments for spawning, juvenile rearing habitat or access to the upper catchment (Arthur & Gray, 2022; Eldon & Greager, 1983). The river also supports several threatened bird species including tarāpuka/ black-billed gull (*Chroicocephalus bulleri*), black-fronted tern/ tarapirohe (*Chlidonias albostriatus*) and over 70% of the total population of wrybill/ ngutu pare (*Anarhynchus frontalis*). All of these species are heavily reliant on braided river habitat (Forest & Bird, 2016; O’Donnell, 2000).

As with many NZ rivers, exotic sportfish species were introduced to create recreational fisheries that have a range of ecological consequences for native species (Jellyman et al., 2017; MacNeil et al., 2024; McDowall, 1990, 2006; Townsend, 1996). These introductions were originally part of a larger ‘acclimatisation’ movement aimed at recreating aspects of Britain under the auspices of European colonisation (Veblen & Stewart, 1982; Williams & Cameron, 2006). The primary sportfish species are brown trout (*Salmo trutta*) and rainbow trout (*Oncorhynchus mykiss*) that each now have a wide distribution (Jellyman et al., 2017; McIntosh, 2000). Chinook salmon (*O. tshawytscha*) runs were also established from planned introductions and hatchery programmes, primarily on the South Island’s east coast where braided river systems provide suitable habitat (McDowall, 1994; Quinn et al., 2001).

All three of the abovementioned sportfish are present in the Rakaia catchment where they were introduced by the Canterbury Acclimatisation Society and are managed today by its modern-day equivalent, Fish and Game New Zealand (Fish and Game). The considerable interest in tracking fishery stocks and recreational use patterns led to the establishment of licensing and monitoring programmes that are specific to each fishery. For the Canterbury salmon fisheries the monitoring activities include annual postal surveys of license holders and fish counts in spawning streams that have been continued for many decades to produce a comprehensive historical record (West & Goode, 1987). Estimates of the annual run size in the Rakaia are now fluctuating around an all-time low of only ∼1000 fish after a precipitous decline from run sizes of >20000 fish in the mid-1990s (Appendix 1, Figure S1) (Dellaway, 2025).

#### Water abstraction framework

The natural flow regime of the Rakaia has been considerably altered by a complex water abstraction regime that provides water for hydroelectricity generation, stock water and irrigation (Appendix 1, Figure S2). As of February 2025, there were 120 consented water takes each of which (with the exception of very small takes) are required to be metered at 15-minute intervals (Terrink, 2021). These data are used to calculate the total abstraction volume and individual permit conditions that are tied to a flow recorder located at Fighting Hill in the Rakaia Gorge (Figure 1). The water balance includes inputs from the Highbank hydroelectric power station which discharges water from the tail end of an irrigation race that has its intake on the Rangitata River, another major alpine river located to the south (Figure 1). Another hydroelectric power scheme is located at Lake Coleridge in the upper Rakaia catchment. This is a naturally occurring lake that is artificially regulated to store water for hydroelectric power generation. Water is diverted into the lake from the Wilberforce and Harper rivers to support this scheme, and outflows from the lake are returned to the river at a discharge point located ∼15 km upstream of the Rakaia Gorge bridge. Since 2013 the stored water has also been used for irrigation purposes under an arrangement between the power company and irrigators (New Zealand Government, 2013). This has the effect of maximising the water take that was previously permitted but at times not utilised by the hydroelectricity scheme.

The difficulty of accurately estimating or gauging river flows in the highly dynamic riverbed is a notable feature of the Rakaia management context. These difficulties are generally approached through linking the various flow considerations and settings to measurements made at the Fighting Hill gauge located in Rakaia Gorge (Figure 1). Estimating the losses to groundwater from the riverbed between Rakaia Gorge and the river mouth is a key example of these difficulties despite its potential to have significant consequences for actual flows in the lower river. In this case the difference is accounted for using an average loss of 21.4 cumecs as a constant in the Rakaia water balance model (Terrink, 2021).

In 1988, the Rakaia became NZ’s second waterway to be granted a National Water Conservation Order (WCO) following a joint application by four Acclimatisation societies with interests in the catchment (New Zealand Government, 1988). The Rakaia WCO and subsequent amendments in 2011 and 2013 play a central and significant role in the policy context for water resource management. It is specifically designed to recognise and protect the outstanding natural characteristics and features of the river (Table 1). Among the key provisions for achieving this is a minimum flow regime that is used by the Canterbury Regional Council (Environment Canterbury) to guide decisions on water abstraction and associated planning mechanisms (Figure 1).

## 2.2 Study design

The study design followed a purposive sampling strategy (Nyimbili & Nyimbili, 2024; Palinkas et al., 2015) that sought the participation of the Rakaia salmon and trout fishing community. To meet the key study objective of capturing the longest standing oral histories from the most experienced members of the community the participant selection criteria included a nominal minimum of 20 years of fishing experience and the additional qualification that it must be specific to the Rakaia catchment. The stakeholder holder engagement methods were tailored towards identifying and engaging with these community members as was reflected in the fishing experience demographic (ranging from 25–75 years’ experience in the catchment).

A target sample of 30 participants was considered appropriate for achieving thematic saturation while also ensuring good geographical coverage of the catchment. In previous evaluations of suitable sample sizes for interview-based studies, Young & Casey (2018) found that 73% of codes were identified in the first six interviews and that substantial theme completion was achieved with sample sizes of 7–10 interviews. Squire et al. (2024) reported that near saturation of in-depth research themes required 15–23 interviews in comparable studies. In our case we were interested in going beyond code saturation to understand contextual aspects of meaning including place-based differences. In an evaluation of similar needs Hennink et al. (2017) found that 16–24 interviews were needed to obtain a full appreciation of contextual meaning whereas only nine interview were needed to generate the majority of all codes in the same study.

Despite the relatively ambitious sample size that was selected for this study it represented a feasible target in consideration of the estimated number of veteran fishers who were thought to be associated with the Rakaia River based on indicators such as the size of the active fishing community at the Rakaia Huts and other popular locations. Project components were established with the funders by way of a terms of reference, meeting minutes (6 May 2024), and discussions at NZSAA’s Annual General Meetings open to all members (2024, 2025). The study was initially promoted through advertisements in fishing community newsletters and presentations. This was followed by a snowball sampling approach in which participants were asked to identify other potential participants who met the selection criteria and could add value to the study (Goodman, 1961). This combination of public promotion and word of mouth recruitment activities was supported by a phone and email contact point through which interested people could register their interest in participating or obtain additional information about the study at any time during the study period of August–December 2024.

## 2.3 Interview process and data analysis

At the commencement of the study the individuals who had expressed interest were contacted individually by the researchers using the contact details and preferences they had provided (phone, email or video call). Following the participant selection criteria, individuals who were more elderly or otherwise known to have a long history of association with the river were prioritised for invitation to join the study. In this initial contact the researcher provided details of the aims of the study, research approach and interview process. Those who indicated a willingness to participate or obtain further information were then sent an information sheet and participant consent form. Interview times and formats were then established to suit the preferences of confirmed participants.

Following the sampling design a total of 30 participants were interviewed between 30 August and 31 December 2024. Qualitative data that were sought in the study were primarily the interview transcripts augmented by visual or audio-visual records that were identified or provided by participants (e.g., photographs, new articles) and in some cases, catch diaries. Interview locations ranged from Timaru to Christchurch. Most were conducted at participants’ homes or at the fishing hut communities located near the river mouth lagoon (Figure 1). Each interview was recorded with the consent of participant which was granted in all cases. Interviews ranged from 30–70 minutes in length and were transcribed within 7 days of the interview. While the oral history approach and use of open questions enabled participants to share information on a wide range of topics, it was important to direct interviews towards the focus of the research questions. This was achieved through a semi-structured interview approach (Yin, 2017) that was based around five groups of research questions guided by a run sheet and set of prompts to be used in the interview (see Supplementary Material). The five research themes were 1) Angler’s connection to the river, 2) Observed change in the environment, 3) Observed change in community, 4) Perceptions of river health and management, and 5) Aspirations for the future.

All interviews were individually transcribed prior to qualitative analysis using an initial round of inductive thematic coding (Boyatzis, 1998), following standard social science methods (Esterberg, 2002; Miles et al., 2013). To ensure anonymity, participants’ names and identifying features were removed from the transcripts and subsequent data analysis steps. Recordings were destroyed following the completion of the project as outlined in the information sheet. All aspects of the study, including participant recruitment, data collection and analysis were subject to an ethics review and approval process following a similar format to that used in previous interview-based studies of fishers in the Canterbury region (Jellyman & Mayall-Nahi, 2022; Rankin et al., 2021) and a previous student research project with salmon fishers in the neighbouring Rangitata river catchment (University of Canterbury Human Ethics approval HEC 2020/95/LR).

## 3. Observations of change

### 3.1 Oral history dataset

The research approach was generally effective in engaging with veteran fishers who have long standing associations with the river as demonstrated by the demographics of the participant cohort (Figure 2A). Most of the participants (73%) were over 65 years of age at the time of the study and the first-hand experience of the river within this age group averaged 53 years. The dataset that was obtained from the study represents 1510 years of first-hand fishing experience that is specific to the Rakaia over a time frame of seven decades. Several participants were the children of fishers who were active in the enhancement the salmon fishery in the 1960s. The combination of their observations and information received from their parents is irreplaceable in the historical record

**Figure 2.**
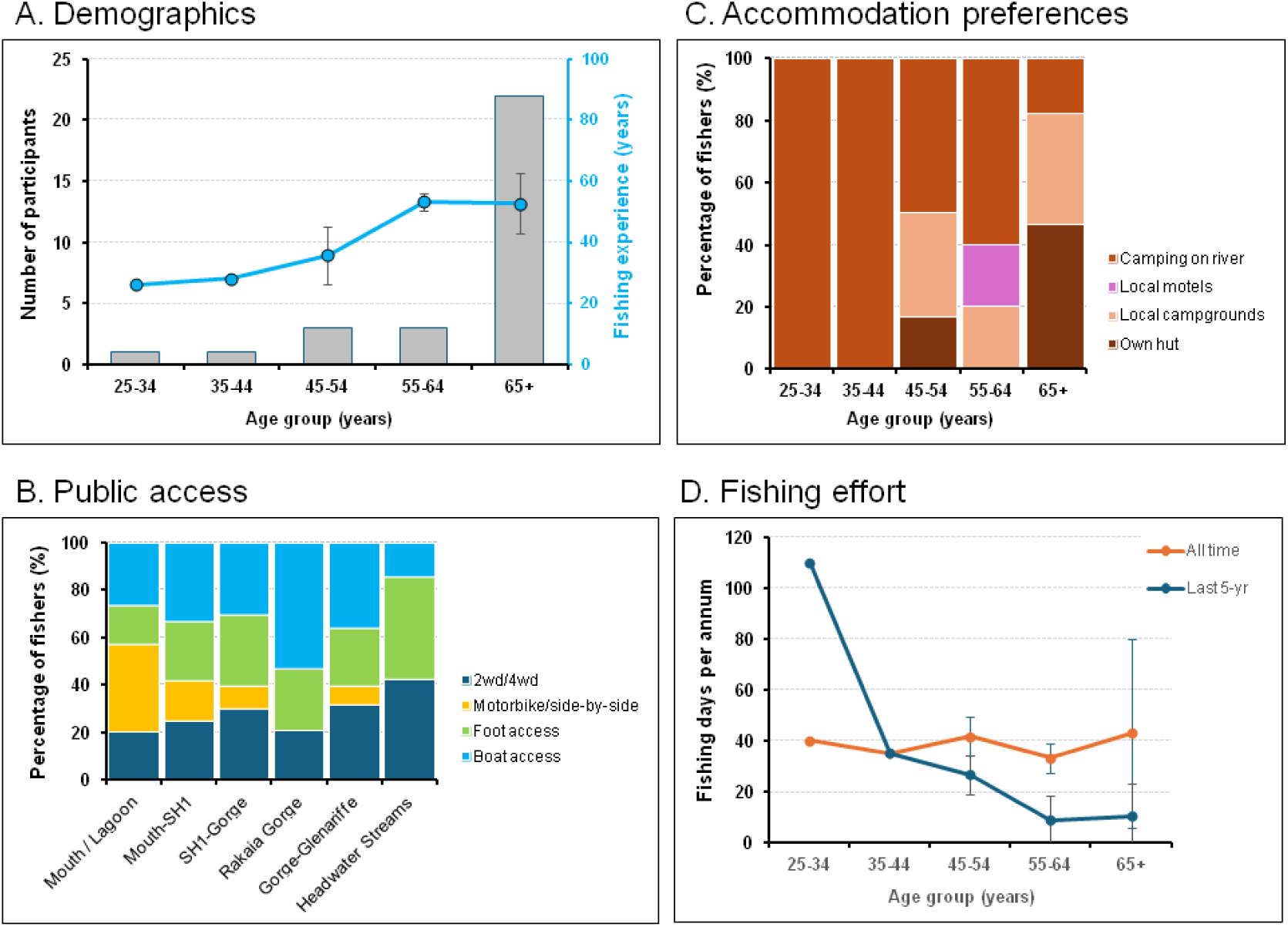
Demographics of the study cohort (n=30). A. Age group and fishing experience of participants. B. Estimated number of fishing days per annum over all time and the last five years. C. Access modes used by fishers in each of six distinct parts of the catchment. D. Accommodation used by fishers.

The locations used for fishing vary across the catchment and traverse several distinct freshwater and coastal interface environments including the river mouth lagoon, braided reaches, Rakaia Gorge, and numerous side streams or confluence areas. Modes of access used by the fishing community include a mix of foot access from road ends, motorbike or off-road vehicle travel on the riverbed, and boat access (Figure 2B). Access by boat access is prominent in the Rakaia Gorge section and at the river mouth lagoon but also includes the use of jetboats to access braided sections elsewhere. Accommodation for overnight stays includes the ownership of fishing huts in the North and South Rakaia fishing communities, particularly in the oldest age group (65+ years), with a trend towards other options in younger age groups (Figure 2C).

Across all participants the estimated annual fishing effort (expressed as ‘fishing days per annum’) averaged 42 days and showed a significant decline to only 16 days per annum over the past 5-year period. The reduction in time allocated to fishing in recent years was highest in the oldest age group who reported a four-fold reduction in their fishing effort in comparison to previous years (Figure 2D).

### 3.2 River environment changes

Geographical changes were mapped across >200 km of the Rakaia mainstem, North Rakaia and headwater side streams. Location-specific data were obtained for over 60 discrete localities that include river reaches described by prominent landmarks (e.g., Rakaia Gorge) and point features such as springs (Figure 3, Supplementary Material Table S1, Table S2).

**Figure 3.**
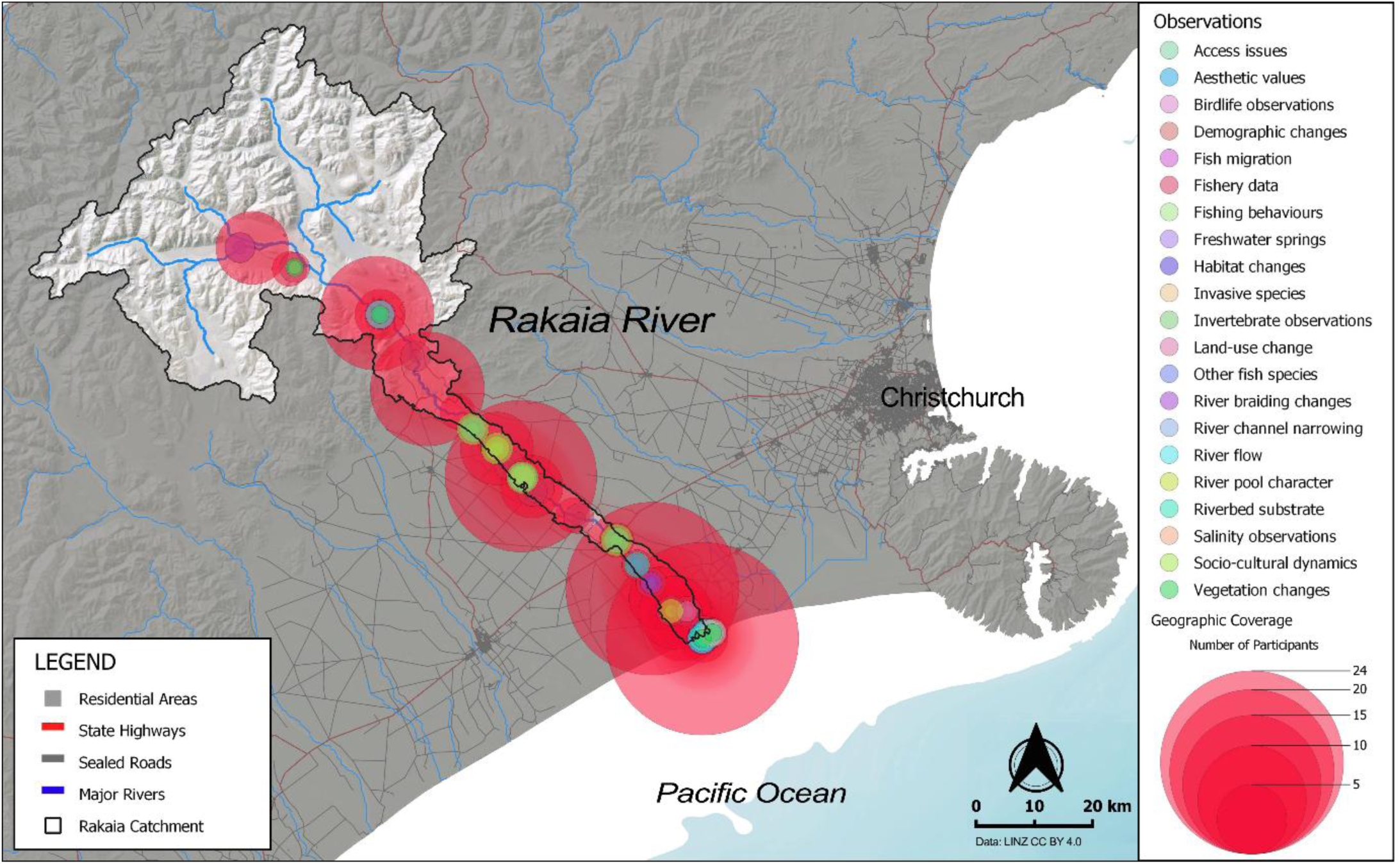
River environment and fishery changes in the Rakaia catchment as recorded by the study participants representing a total of 256 discrete geographic observations.

#### River features

Changes to the physical environment involve a wide variety of natural features and river attributes including riverbed character, substrate size, flow patterns, water clarity and riparian environments (Figure 4A, Supplementary Material Table S1). For many anglers these changes were associated with a reduction in suitable fishing pools and runs, particularly below Rakaia gorge. For example: “When I first was fishing the Rakaia, particularly from Sleemans Road down there would be holes around the river. Now you are going further and further to find the holes, and they are nowhere near as good as what they used to be” [P20]. Many observations related to changes in sediments and particle sizes within the river system, especially a reduction in the size of stones and boulders below the gorge and at the river mouth. This idea was captured by one participant who noted “So now, if you walk along the spit it is like walking on ball bearings you know, but on the inside … it was very, very large boulders, quite rocky” [P9]. Observations recorded by many participants included an increase in siltation in the mid reaches below SH1, an increase in riverbed elevation, and a reduction in the height and extent of the gravel bank adjacent to the river lagoon.

**Figure 4.**
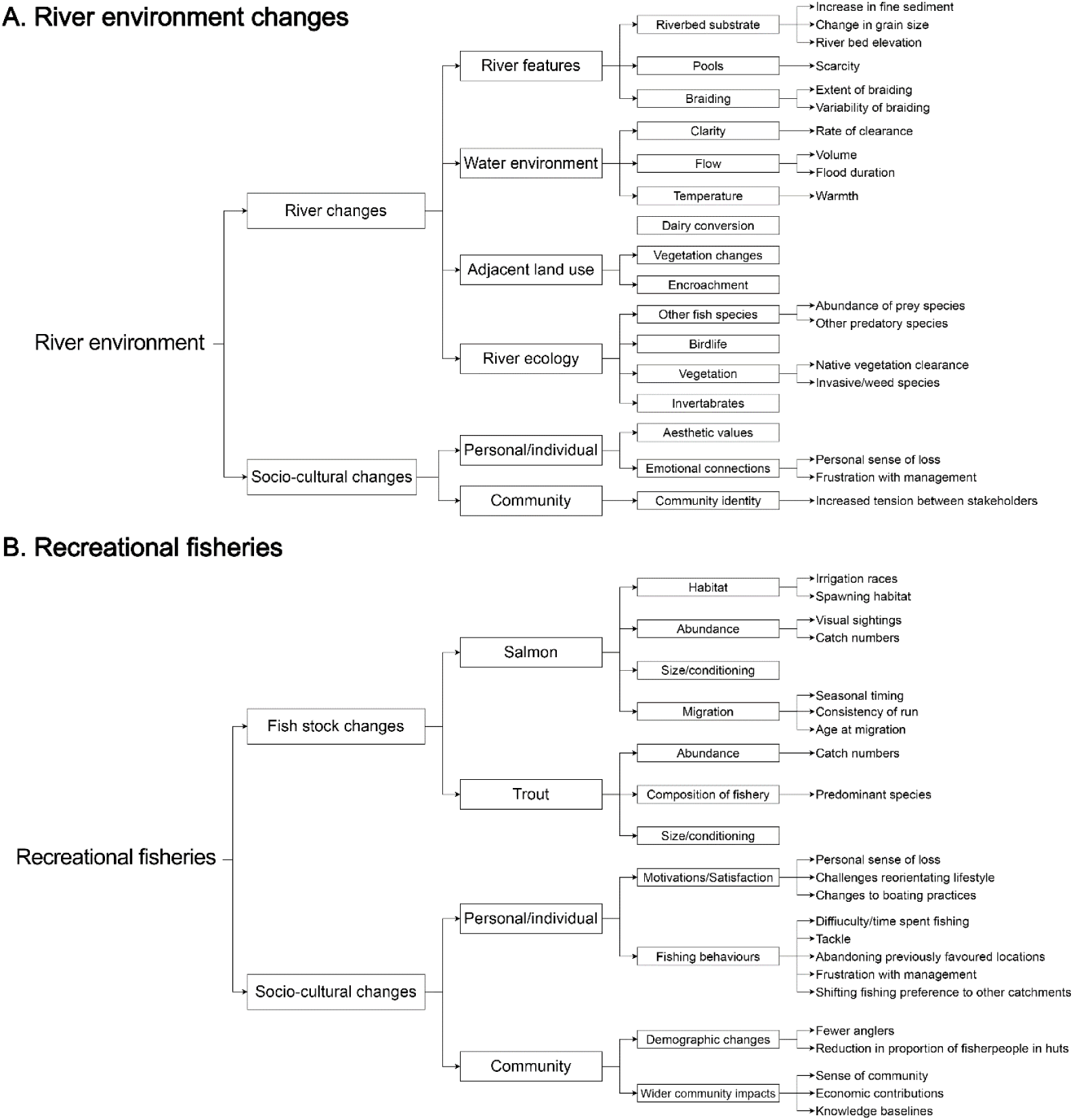
Results from thematic analysis of oral histories of recreational fishers (n=30) in the Rakaia catchment divided according to two overarching sets of questions A. River environment changes. B. Social-ecological changes in recreational trout and salmon fisheries.

Changes recorded in and around the river mouth area include a narrowing and shrinking of the lagoon, an increase in susceptibility to flooding and the disappearance of freshwater springs. One such spring was described as follows: “There was another massive spring halfway up the lagoon in the centre … there used to be a bulge in the water in the lagoon that was how much water was coming up. It was massively deep. I used to try to go through it with a wetsuit, it used to just, I used to end up at the surface in a hurry” [P17]. Several participants noted public access related changes including the loss of access through land-use change and challenges in launching boats or utilising areas of the river which had become less suitable due to shallower water depths in those reaches. Physical barriers generated by water intake structures were also described: “Because of the way they do their irrigation intakes now, far above where the actual intake is you can’t cross … so you can’t get to the river” [P28].

#### Water environment

Fifty observations regarding changes to water were made by anglers in this study including observations relating to clarity, flow level, water quality, temperature and water outtakes. Many participants reported a reduction in the overall river flow: “The river certainly has changed there is no doubt about that, it is nowhere near what it used to be, it is nowhere near the volume of water” [P9]. Additionally, participants noted alterations to the ways in which water volumes rose and fell. The most notable trend involved river floods receding more rapidly than in the past with rapid drops in water volume becoming more commonplace. Participants related this knowledge and understanding of flow conditions to their influence on conditions which were suitable for fishing. Observed changes include: “In the 70s, 80s and 90s when the river came down in a flood from the norwesters and the snow melt that would take days and days to clear because there was a lot more water. Now it just takes 2 to 4 days, depending on the size of the flood, to clear” [P30]. These changes were also connected with the characteristics and duration of flow ranges that supported successful fishing. For example: When I was young … we would start to look at going fishing when the river was getting around 200 to 220 cumecs, that would be just about fishable. Nowadays … I would start looking around at 120” [P27]. Several participants also noted that the water temperature in the river is generally higher than it used to be. For example: “The temperature is higher… you can tell straight away as you’re walking through the river what the temperature is” [P1].

At the river mouth the reduced flows were equated with an observed inability of river flow to slow incoming tides leading to earlier closures of the river mouth and more extensive saline intrusion. These alterations in water volume and flow rate were also associated with increased difficulty in navigating the river in a jet boat, a decrease in anchor size required for fishing off the mouth, and the need to use lighter fishing tackle to suit the flow of the river. One participant noted: “In the early days up to the 2000s because of the sheer volume of water in the river the fishing tackle was minimum of 10kg line and the hook size was 28 grams which is a reasonably heavy spinner. Now I use, 6 or 7kg line with a very small spinner only maybe 15 grams and a very small hook because you don’t need the weight of line and spinner to get down to where the fish are” [P10].

#### River ecology

Many participants noted the dramatic decline in observations of Stokell’s smelt in the lower river consistent with previous reports. This reduction was described as “… beyond decline … they used to come in like a big run of whitebait on the coast, it would be about a metre wide and they would go all day” [P5]. Other observations related to fish species included recent observations of haku / kingfish (*Seriola lalandi*) at the river mouth, decline in the whitebait catch, and no longer observing squat lobster (*Munida gregaria*) offshore from the river mouth. Participants also reported declines in characteristic birdlife, notably a reduction in the tern and gull colonies that are a defining feature of the river ecology. The observations included reports of birds in poor condition that were thought to be associated with starvation events or other hardships.

### 3.3 Recreational fisheries

The thematic analysis identified over 30 discrete social-ecological aspects of the recreational fishery changes (Supplementary Material Table S2). Three major groups of themes collectively describe these effects: the status of the fish species, participation in fishing, and associated impacts on local communities (Figure 4B).

#### Fishing success

All participants described a significant decline in fundamental indicators of the salmon and trout fisheries over the course of their association with the river including familiarity with the much-reduced run sizes as estimated from Fish and Game surveys (Appendix 1, Figure S1). Information on fishing success in the salmon fishery generally reflects these trends with many participants noting that the fishery had deteriorated markedly since the mid-1990s. In addition to reduced catches, several participants described a marked reduction in observations or sightings of salmon rising or travelling up the Rakaia River. These reports contrasted significantly with earlier experiences of fishing on the river, with one angler noting “I can remember counting over 100 salmon shoot a bar in a day, two and three at once, beautiful to watch, amazing” [P29]. Catch per unit effort was widely regarded to be a critical issue. For example, one participant noted ‘If I caught one or even two a season I would be lucky, before that, I mean you go down and I would catch a couple a weekend, there seemed to be plenty there” [P6].

These circumstances have influenced the reduction in fishing effort in previous years in older age groups (Figure 2D) and reflect a general sense of despondence in the fishing community. In the words of one participant: “it’s absolutely gutting to see my hobby slowly disappearing or slowly coming to an end.” [P15]. Another noted how the decline in fish size meant “it is very hard to get motivated with a 4-pound fish and that is coming from an avid fisherman.” [P27]. For many the severity of the decline undermined their ability to engage with what was previously a primary passion or source of recreation. For example, one angler described that “it was a massive part of my life salmon fishing and it’s just gone it’s just absolutely gone.” [P8]. This was strongly felt among anglers who had anticipated increasing their fishing activities in retirement. In response, many participants reported shifting their expectations of fishing success. One angler described related social effects: “… if you were going out to a party and someone or someone rung you and wanted you to catch them a salmon if they were having a wedding or something you could just go down and catch one, but now you’re chances of catching one would be that slim it’s not funny.” [P17]. Another noted: “when I talk to other blokes … it’s basically, have you seen anything? Not, have you caught anything?” [P7].

#### Participation and personal wellbeing

Many participants noted a marked decrease in the number of people fishing on the Rakaia, particularly at the river mouth. For example: “when I started fishing down there it was like picket fence material at the mouth. It was tangles and god knows what … 70–80 people. You don’t see that anymore.” [P14]. Another angler noted similar changes stating: “When I started there would be a hundred people fishing sort of thing, now there is only five a day.” [P2]. Many participants also noted a generational shift in the demographics of the fishing community with very few new anglers developing an interest in fishing the river. One participant stated: “There does not appear to be any younger ones there now, really keen young ones, there are some don’t get me wrong but not like they used to be.” [P7]. Additionally, these observations of reduced participation were accompanied by accounts of fishers who have abandoned the Rakaia in search of other catchments thereby placing additional pressure on other fisheries.

Participants reported significant emotional and psychological consequences from their experience of changes to the fisheries and river environment. The words of one angler were echoed by many: “I think it is hugely depressing, emotionally depressing.” [P23]. For many participants the widespread changes they had witnessed had an impact on their sense of place in the river. One participant observed that: “There’s nothing left that’s wild anymore … once upon a time you wouldn’t see or hear any farming activity, now you go up here and there is a damn fence right along the edge of the river.” [P20]. Others perceived that the changes had materially disrupted their experience and connection with the river. In the words of one participant: “It is like having a death in the family, it’s like a dislocation and a lot of it doesn’t need to happen. It affects me personally, I get very angry about it…” [P22].

#### Community impacts and identity

Many participants noted that this trend of anglers leaving the North Rakaia Huts had resulted in a widespread change in the sense of community in the area as the demographics shifted from a once predominately angling based population to one of holiday makers and permanent non-angling residents. One participant recalled: “When I first started going everyone was there for one reason, fishing, and everyone fished. Now … they are just there because it suited their price range and a house and it’s an easy trip into town.” [P28]. The declining fish catches have had an impact on the sense of community associated with the river and its fisheries. This was described by one participant: “People were absolutely passionate about the river and the fishing, there were big fish catches, big numbers of fish being caught, so they were the heydays.” [P26]. Similar effects were also noted in other towns where salmon angling had once been an important part of local culture. One participant described: “… you would see people with rods hanging out of their car for the salmon season, well nowadays you would never see anyone with a rod hanging out their car, the whole dynamic of the salmon fishery has changed. Something has gone. In my lifetime it’s just about disappeared. [P23].

Additional community and socioeconomic dimensions include declining opportunities that have prompted some members of community to either give up angling or to leave the area altogether. One participant described their reasons for reallocating previous investments in the area: “Put it this way … I’ve just sold the bach (fishing hut). That’s where I’m at, at the moment. I’ve basically walked away from it …” [P2]. Participants also noted the increased difficulty to sustain angling related businesses and the unlikeliness of re-establishing an angling guiding business on the Rakaia River. For example: “If I wanted to restart my guiding career, I don’t think I could do it now. I certainly wouldn’t be able to offer the same level of fishing for them.” [P13]. The lack of new engagement with the fisheries also raised concerns for the future knowledge of the fishery and community advocacy for its resources. For example: “I get a little scared we’ve got these guys that are pushing the 60s and they are all guys that I look up to, when they are gone are we going to have this younger generation sticking up for our rivers and sticking up for our fisheries? Where is the next generation coming through?” [P27].

## 4. Discussion

### 4.1 Utility of local ecological knowledge

This study presents an example of participatory research with community members who have long histories of accumulated local knowledge that originates from a high degree of place attachment. In the Rakaia fishing communities this arises from a high number of repeat visits to the same locations over considerable time periods and is further reinforced by cultural developments such as the establishment of fishing hut communities (aggregations of small cribs and overnight accommodations). This example also has many parallels with other salmon fisheries globally where salmon runs have deep cultural and traditional values and significance for local communities and Indigenous peoples, particularly in North America (Atlas et al., 2020; Carothers et al., 2021).

Previous studies have shown that detailed local knowledge can develop from long-standing associations with specific features and places (Chapin & Knapp, 2015) and that this process facilitates the identification and understanding of change in the landscapes and seascapes (Campbell & Orchard, 2023). Our focus on the collective knowledge of highly experienced resource users is likely transferable to many other contexts where there are similar longstanding and localised relationships between natural resources and resource users (Kitolelei et al., 2022; Mixon & Caudill, 2017; Ruddle, 1996). These relationships are frequently encountered in place-based activities such as fishing, wild harvest and outdoor pursuits where participants are reliant on and attuned to specific environmental features and conditions (Orchard, 2025; Veitayaki, 2002). Studies of local knowledge in North America have also highlighted the accumulated traditional knowledge and associated cultural significance that is linked to specific resources such as seaweed (Gitga’a) and salmon (Tlingit) (Menzies, 2006). In a wider historical context, the development of local ecological knowledge in Indigenous cultures often arises from environmental feedback on traditional practices for natural resource management that are passed down to successive generations (Berkes et al., 2000). From this perspective the accumulation and transmission of collective knowledge may also be motivated by benefits to individuals and communities. For example, García-Quijano (2009) found that local ecological knowledge in small-scale Puerto Rican fishing communities was driven by its positive influence on fishing success in the dynamic and rapidly changing coastal environment.

While there is a growing recognition of the utility of social data for resolving trade-offs and supporting conservation planning initiatives (Knight et al., 2010; Whitehead et al., 2014) the use of community-held data as a source of scientific knowledge remains a largely untapped resource in Western cultures. However, participative mapping and volunteered geographical information (VGI) techniques have long been recognised and utilised in community-based natural resource planning in developing countries (Chambers, 1994). New technologies are also facilitating an increasingly diverse range of community and citizen science initiatives to support public involvement with science (Coulson et al., 2021; Newman et al., 2012; Orchard, 2019; Palacin et al., 2020). Oral history research further engages with these opportunities as a form of participatory science with knowledge holders (Hutching, 1993). As a people-centric methodology it offers an inclusive and broadly applicable approach that is well aligned with local and traditional knowledge systems based around oral traditions and storytelling (Portelli, 1992).

As shown in this study, accumulated local knowledge may be particularly useful where it can help to illuminate past conditions that remain important for contemporary decision-making because they represent management baselines or potential reference states for environmental restoration. Moreover, the historical period that predates the widespread availability of high-resolution satellite data products (i.e., pre-2000s) is of particular interest due to the much-reduced opportunities for retrospective remote sensing in comparison to modern sensors and datasets. As is common with community science approaches, questions around the validation of information from individual observers has often hindered its recognition in formal decision making (Ottinger, 2010). However, collective knowledge analyses can facilitate the triangulation and corroboration of discrete observations and build evidence for common themes and trends across observers (Peach et al., 2021; Romney et al., 1986).

### 4.2 Social-ecological interactions

Several notable linkages between changes in the river environment and associated socio-cultural values were identified in this study. These transcend beyond the personal reward and wellbeing values of fishers in the recreational fisheries, to include impacts on livelihoods, community amenities, identity, social interactions, demographics and property ownership patterns (Figure 5).

**Figure 5.**
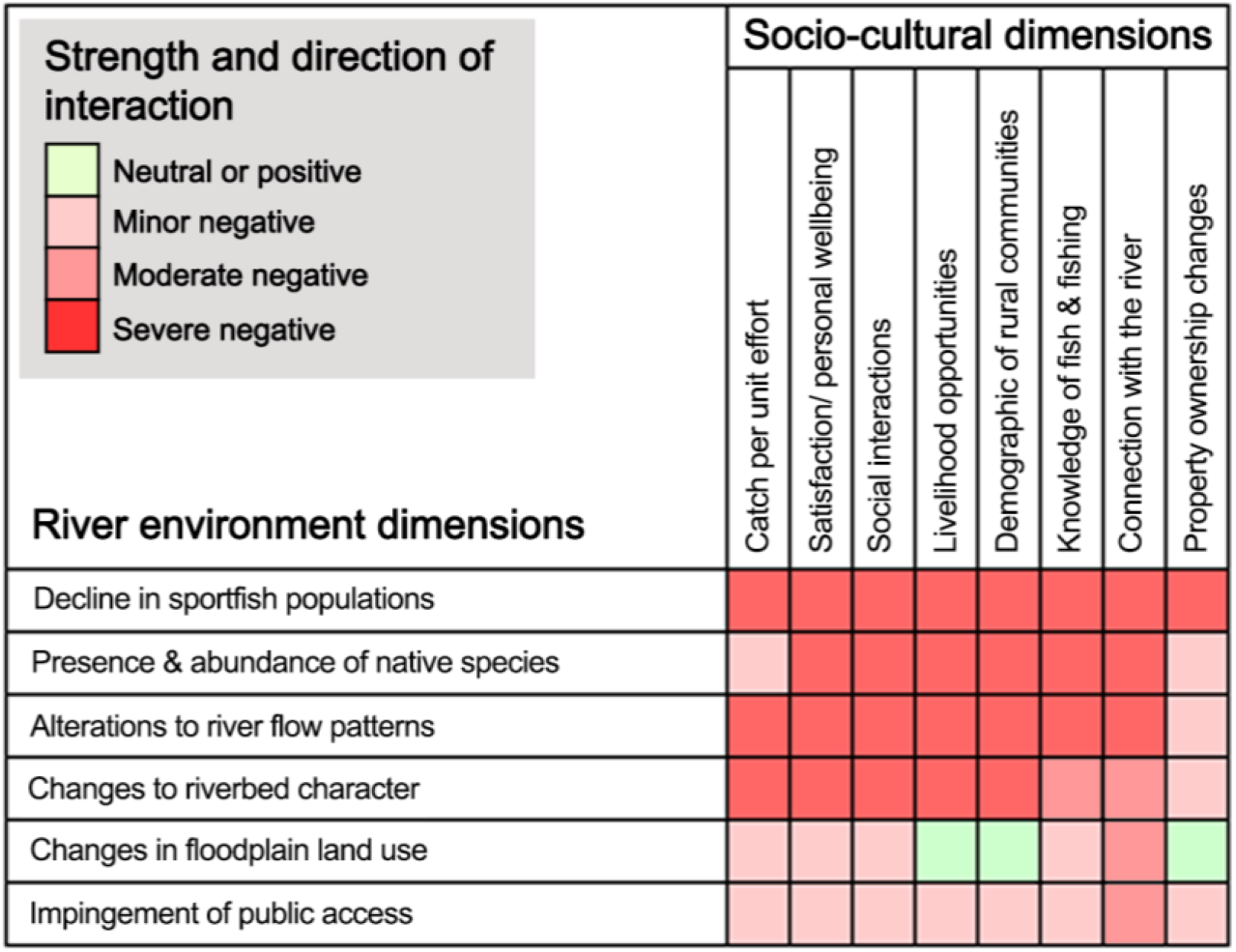
Linkages between the natural features and ecology of the river environment and the socio-cultural values of fishing communities in the Rakaia catchment.

Undesirable social impacts that are linked to the declining fisheries include despondence in the fishing community, reduced participation or relocation of fishing effort, abandonment of fishing hut communities, and generational effects such as the loss of knowledge associated with fishing and reduced connection between people and the river. Changes in presence and abundance of native species also had negative effects on socio-cultural values that were linked to perceptions around their role as indicators of wider ecosystem decline. Physical changes to the riverbed character have many direct and indirect effects on socio-cultural values. Many of these are associated with changes to the natural features that are used for fishing and other outdoor activities but also include a notable interaction with public access to the riverbed. Similarly, the river flow regime has a direct effect on nearly all socio-cultural values through its influence on many other environmental components (Figure 5). Examples include the influence of low flows on the extent of instream habitat, nourishment of riparian wetlands, sediment flushing and settlement regimes, river bar barrier behaviour, and extent of the freshwater plume at the river mouth as a brackish environment with important connectivity functions (Devlin et al., 2001; Gray et al., 2018; Measures et al., 2020; Orchard et al., 2025; Sheldon, 2017).

The concept of amenity values is defined under the Resource Management Act 1991 (RMA) as the natural or physical qualities and characteristics that contribute to the cultural and recreational attributes of an area (New Zealand Government, 2025). Examples in the Rakaia catchment include open water features and clear water environments at the river mouth lagoon that are reported to have declined in extent or condition over the timeline of this study. These changes affect amenity values for their associated communities whilst also representing habitat and ecosystem level changes.

Other significant findings include the loss of naturally occurring springs in the lower river and evidence that the Rakaia River North Branch is undergoing a hysteresis or ‘flipped state’ from a previously clearer water state. The regional context already includes a well-known example of a persistent flipped state in Te Waihora / Lake Ellesmere, a nearby coastal lagoon that was previously a clear water environment dominated by extensive aquatic plant forests (e.g., Ruppia spp.) (Hughey et al., 2013; Hughey & Taylor, 2009). Its history includes the demise of a previously world-renowned brown trout fishery and decline of other culturally important species (Hardy, 1989). Despite decades of management attention these historical circumstances remain a concern and are currently the focus of an extensive cultural, ecological and educational programme that is seeking solutions to restore the lagoon for future generations (Te Waihora Co-Governance Group, 2025). The establishment of undesirable stable states as exemplified in this example is a thorny issue for natural resource management due to their potential to invoke loss or damage to existing resources and relative permanence (Beisner et al., 2003; Nyström et al., 2000; Scheffer et al., 2001). As we highlight for the Rakaia North Branch environment, the most effective strategies require an understanding of tipping points beyond which a desirable state cannot persist as a focus for management actions (Mushet et al., 2020; Suding & Hobbs, 2009). In this case the hydrological processes are influenced by both the connectivity of overland flow paths and the location of freshwater springs in the lower catchment.

### 4.3 Implications for river management

Across the local knowledge documented in this study, over 20 discrete components of environmental change can be identified that directly affect the outstanding river characteristics and features that are protected under the WCO (Figure 6). This is significant since the WCO sets out the relevant objectives for these characteristics together with clear guidance on the baseline states and time frames for compliance and effectiveness evaluation (New Zealand Government, 1988). As such our results imply a need to at least investigate each of these changes to determine the degree to which they are influenced by anthropogenic factors, and may therefore be manageable, or alternatively should be regarded as a natural component of the river’s dynamics and therefore consistent with the WCO and other relevant policy objectives. Key opportunities for improving the management of trends that are signaled in these findings include an improved understanding of changes to natural features such as pools, holes and springs, the extent and condition of the river mouth hāpua, vegetation patterns in riverbed, riparian and aquatic environments, and interactions with land use and public access. These aspects are additional to the more complex questions involving the drivers of decline in wildlife and fisheries values and should be regarded as essential preliminary investigations that may help to develop regenerative approaches to halt decline and promote restoration and resilience.

**Figure 6.**
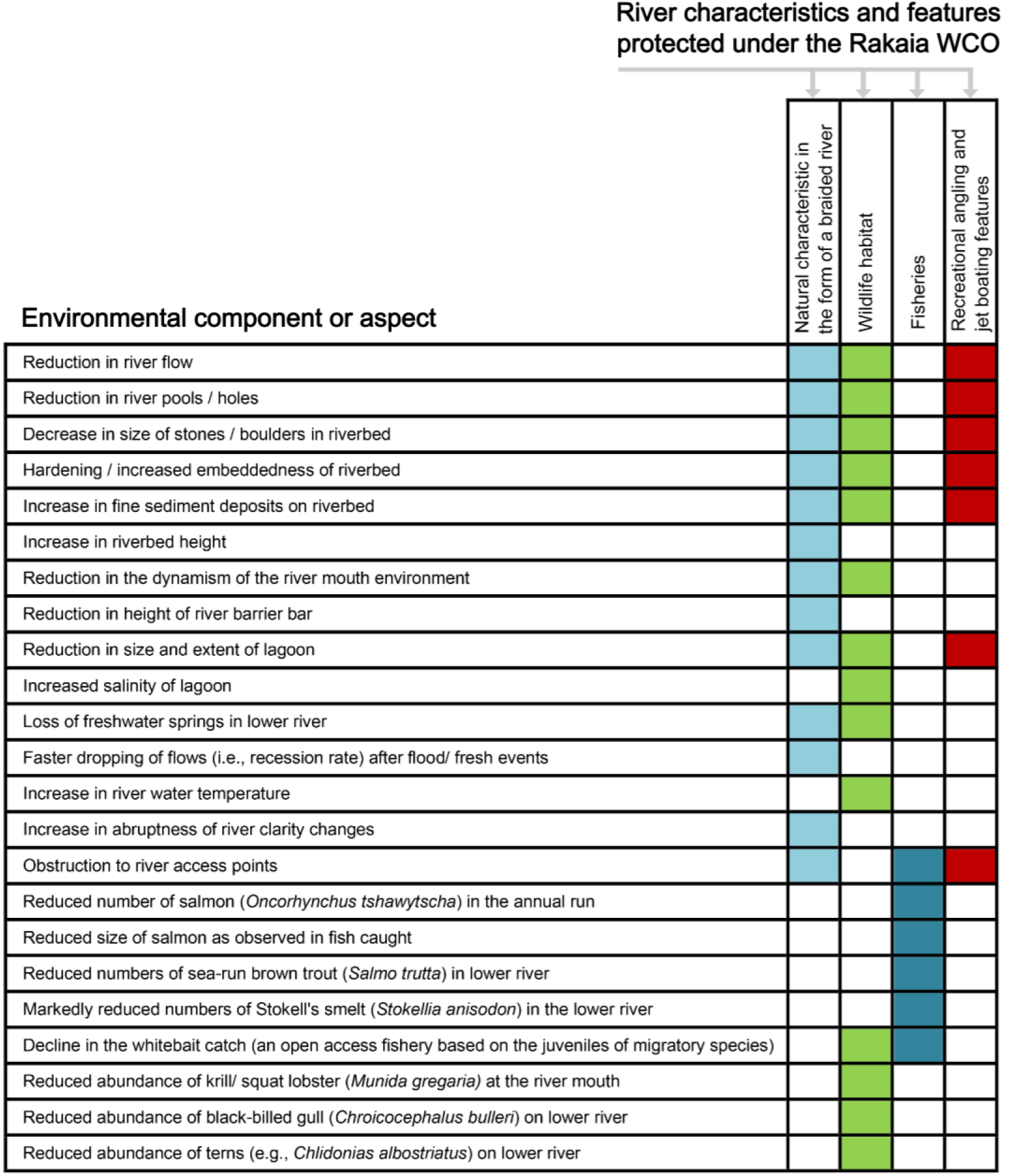
River environment, fishery and ecology changes observed by fishers in the Rakaia River catchment mapped against the requirements of the National Water Conservation (Rakaia River) Order 1988 (WCO) (New Zealand Government, 1988).

To date, only a few of these topics have been the subject of management attention with the notable exception of recent investigation into the drivers of Stockell’s smelt decline (Arthur & Gray, 2022; Hickford, 2022; Jellyman & Mayall-Nahi, 2022) and earlier work by the Canterbury Regional Council (Environment Canterbury) on the extent of land-use change on braided riverbeds in the region (Gray et al., 2018; Grove et al., 2015). There is also currently no comprehensive management or monitoring plan at the Rakaia catchment scale. Factors contributing to the policy implementation gap include inertia in the establishment of environmental monitoring in the catchment, together with the lack of any formal evaluation of the WCO against its objectives. A crucial element that is currently being tested in court proceedings concerns the institutional locus of responsibilities for WCO monitoring and enforcement (Ministry for the Environment, 2023). These proceedings have a specific focus on clarifying the role of the regional council as the key government agency responsible for implementing the WCO and concerns around the use of Lake Coleridge as a source of water for irrigation.

On a longer timeline these recent developments are just the latest of many flow-related proposals and administrative processes that have influenced the history of water abstraction from the catchment (Appendix 1, Figure S2). They further illustrate the importance of a firm commitment to adequate monitoring to guide the implementation of environmental policy towards its intended objectives. The detection of decline from historical baselines plays a foundational role that may also be limited by its inherent reliance on the availability of historical information. Participatory approaches with knowledge holders can help to address this information gap and improve the understanding of environmental change over meaningful timelines.

## Appendix 1. Supplementary material

**Figure S1.**
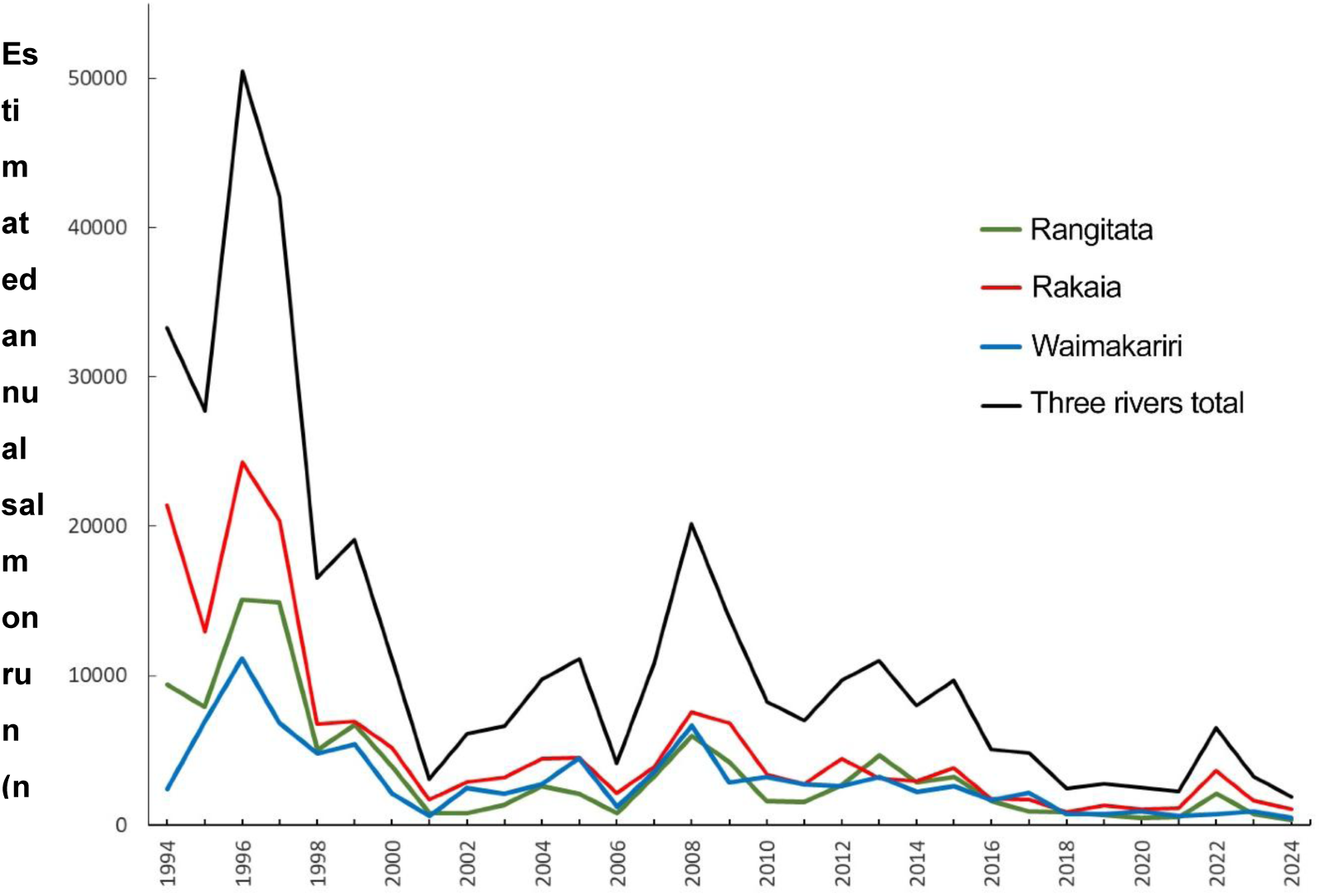
Estimated annual salmon return for three major braided rivers in the Canterbury region (Rakaia, Rangitata and Waimakariri) over the period 1994-2024. Adapted from Dellaway (2025).

**Figure S2.**
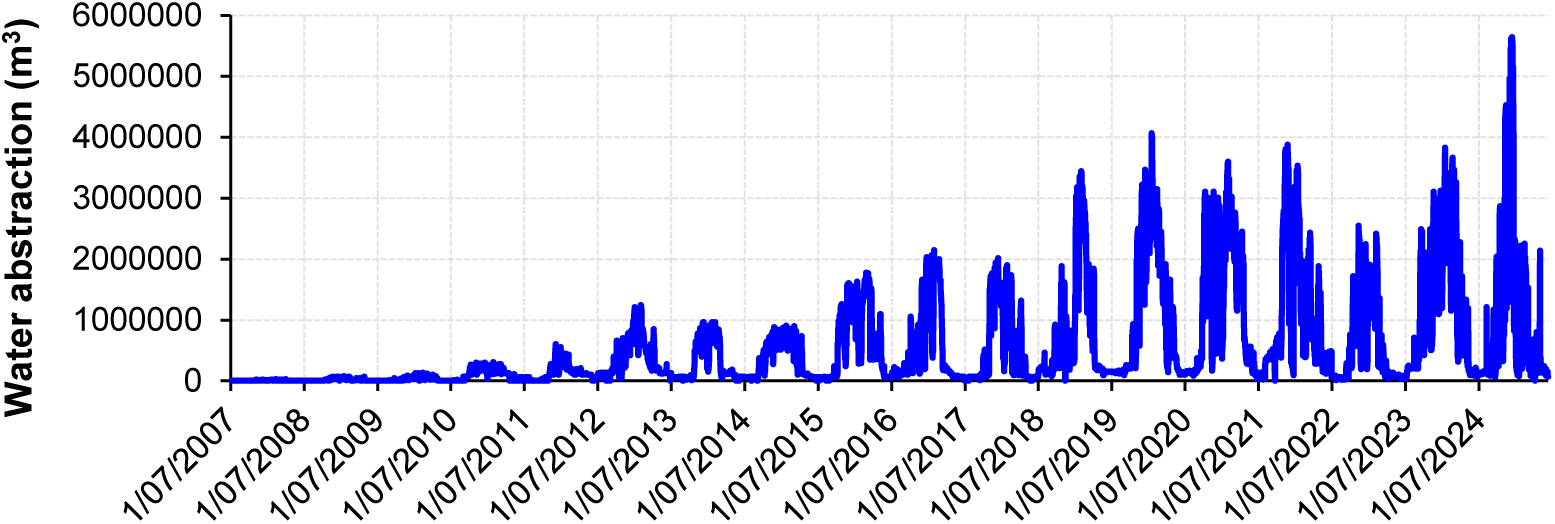
Water abstraction volume in the Rakaia catchment since 2007 as measured from metered water takes. Source data courtesy of Environment Canterbury.

## Data Availability Statement

Dataset available on request from the authors subject to privacy provisions.

## Acknowledgments

We acknowledge and sincerely thank all of the study participants who contributed their time and knowledge to this project. Funding was provided by Future Rivers and the New Zealand Salmon Anglers Association (NZSAA). Logistical support for survey promotion was provided by NZSAA and Fish & Game. Water abstraction data from metered water takes was kindly provided by Environment Canterbury.

