## Supplementary Material for "Combining local knowledge from oral histories and participatory mapping with veteran fishers to identify long-term environmental change in a nationally significant river"

--

#### Graphical abstract

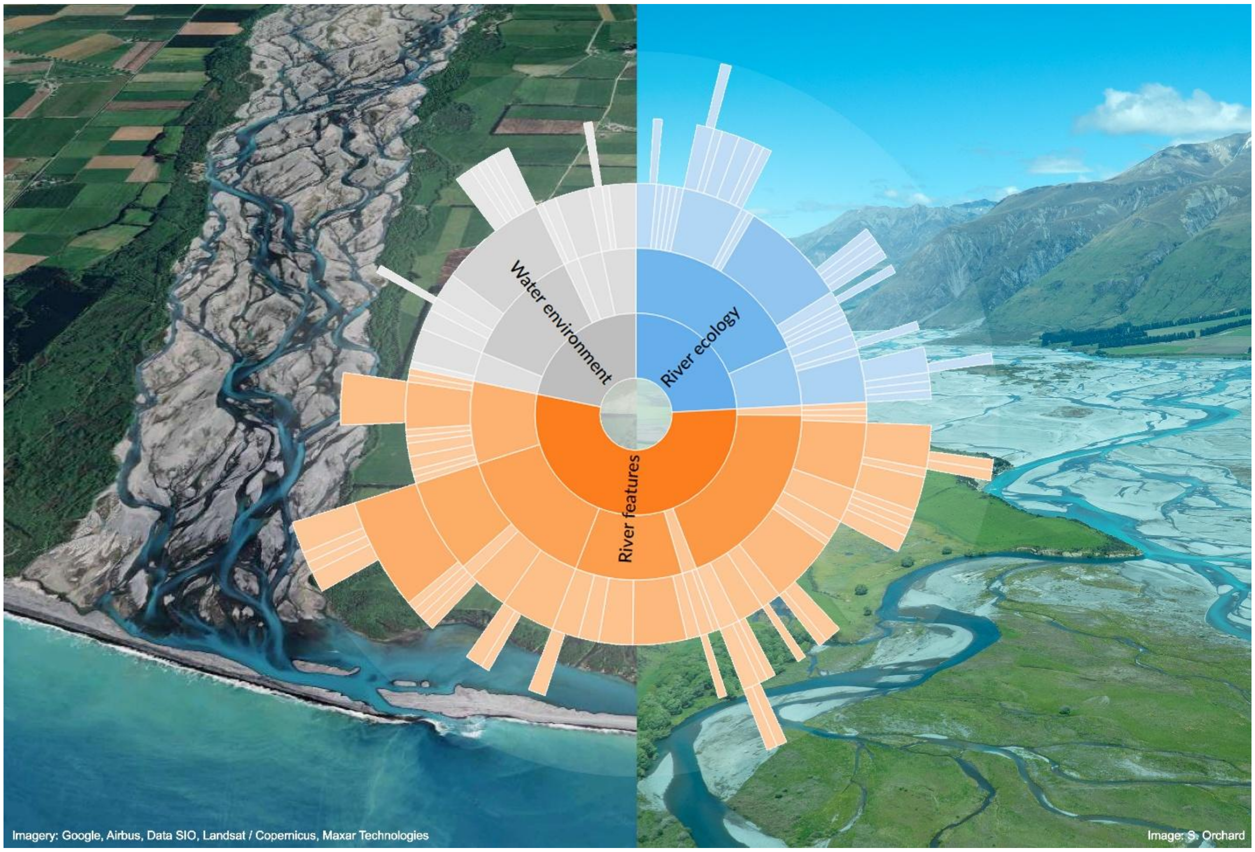

**Table S1**. River environment and ecology changes observed by fishers in the Rakaia River catchment.

| Components | Observed change | Local context |
|---|---|---|
| *Physical environment and river features* | | |
| River pools | Observed reduction in river pools / holes | "The holes and pools that used to be in the river are not there." [P10] |
| Extent of river features and variability. | Observed reduction to in-river variability. | "She was a much more dynamic river. Now the mouth will set a format." [P5] |
| Firmness of riverbed | Observed hardening of riverbed | "I noticed that the riverbed started to get harder." [P22] |
| Size of substrate | Observed decrease in size of stones and boulders across river length. | "The boulders were pretty big in those days, well now they don't exist." [P9] |
| Presence of silt | Observed increase of siltation throughout riverbed. | "Today you pick up a stone it's covered in bloody silt." [P4] |
| Riverbed elevation | Observed increase in riverbed height. | "Shingle is coming down all the time it is just filling up and we haven't got the big flows to get rid of it." [P14] |
| River barrier bar structure | Observed reduction in height of river barrier bar. | "The height of that shingle bank from the time I first knew it to when we left, it was just gone." [P28] |
| Lagoon size and extent | Observed reduction in size and extent of lower lagoon. | "We used to water ski down there in the lagoon and now it is probably as wide as this room here, and you couldn't ever do that now." [P6] |
| Salinity of lagoon water | Observed Increase to salinity of lagoon. | "We had crystal clear water in the lagoon which has now got salt in it." [P17] |
| Presence of freshwater springs | Observed loss of freshwater springs in lower river area. | "You don't see the upwelling coming from the springs down at the mouth anymore." [P22] |
| Flood hazard | Observed increased flood risk to mouth adjacent village. | "We are starting to see it now with more flooding down here." [P17] |
| River flow volume | Observed reduction in primary river flow volume. | "The Rakaia was a massive river and now it is more a shadow of its former self." [P3] |
| River flow regime | Observed increase in rate of flood flow recession. | "It would be six weeks before the river was clear. Now you get a 500 or 600 flood two days later it's down to 150." [P28] |
| River water temperature | Observed increased in river water temperature. | "It's cold but it's not that really icy cold like it used to be." [P30] |
| River water clarity | Observation of increase in rate at which river clarity increases following a fresh. | "This river can go from unfishable to too clear within a window of about 5 or 6 days now. " [P15] |
| Availability of river access points | Observation of obstruction to river access points. | "You can't get to it because of the way they do their irrigation intakes now." [P28] |
| *River and flood plain ecology* | | |
| Whitebait (An open access fishery based on juvenile galaxiid species). | Decline in whitebait catch (An open access fishery based on juvenile galaxiid species). | "The whitebait numbers are not around." [P30] |
| Smelt | Observations of reduced smelt (Stokell's smelt - *Stokellia anisodon)* numbers. | "The smelt are not there, they used to have them, and they'd be there in millions." [P2] |
| Krill/squat lobster | Fewer observations of squat lobster (*Munida gregaria)* at the river mouth. | "You could see acres of it, well that's sort of gone too you know." [P16] |
| Bellbirds | Fewer observations of bellbirds (*Anthornis melanura)* in upper river area. | "It's usually got a number of bellbirds up there but that has diminished." [P1] |
| Terns | Observations of reduced tern numbers in lower river. | "We don't have the tern rookeries we used to have." [P19] |
| Black-billed gulls | Observations of reduced gull (*Chroicocephalus bulleri*) numbers in lower river. | "Black billed gulls ,there would be flocks of hundreds or thousands of them, they are not as prevalent." [P10] |

**Table S2**. Fishing success, participation and fishing community changes in the Rakaia River catchment.

| Dimensions | Observed change | Local context |
|---|---|---|
| Fewer fish | Catch numbers | "*I can remember going there in a morning, in those days and you'd be ten or twelve fishermen fishing it, in a hole, and there'd be ten fish caught. Now you'd go there, and you might see six fishermen, but you probably wouldn't see any fish caught, you might see one."* [P7] *"… quite often we would catch two or three a day if you were on the right spot. And then its declined now, so two a year."* [P20] |
| | Fish observations | *"The last years that we were fishing, or I was fishing, you would rarely see a fish. The fish just weren't there."* [P26] |
| | Decline since mid-1990s | "*The fishery was strong for many years, but about '92 that's when we started to see the real decline and it just sped up. It really sped up and on through the 2000s."* [P18]. |
| Smaller fish | Size of fish landed | *"The fish used to average at 15 pounds, certainly a fish in the teens, now they have got smaller and smaller. Nowadays you are lucky if a good fish is over 10 pounds."* [P11] *"they (trout) are near non-existent compared to when I was a young fella. It would be nothing to get a dozen fish, and they would be over 10 pound now a good sea run is like 5 or 6 pound."* [P24]. |
| Fishing effort | Catch per unit effort (CPUE) | '*If I caught one or even two a season I would be lucky, before that, I mean you go down and I would catch a couple a weekend, there seemed to be plenty there*." [P6]. |
| Participation in fishing | Fisher numbers | *"When I started there would be a hundred people fishing sort of thing, now there is only five a day."* [P2]. |
| | Fisher demographics | *"does not appear to be any younger ones there now, really keen young ones … not like they used to be in our days, and I think it's the fact that people turn around and realise its buggered."* [P7]. |
| Social impact | Participant satisfaction | *"I think it is hugely depressing, emotionally depressing." –[P23].* |
| | Personal wellbeing | *"It is like having a death in the family, it's like a dislocation and a lot of it doesn't need to happen. It affects me personally, I get very angry about it…"* [P20] |
| | Sense of loss | *"It was a massive part of my life salmon fishing and it's just gone it's just absolutely gone."* [P8] |
| | Shifting expectations | *"when I talk to other blokes am I going out with fishing and that, it's basically, have you seen anything? Not, have you caught anything? Have you seen anything?"* [P7] |
| Community impact | Community demographics | "*… you would see people with rods hanging out of their car for the salmon season, well nowadays you would never see anyone with a rod hanging out their car, the whole dynamic of the salmon fishery has changed. Something has gone. In my lifetime it's just about disappeared.* [P23] |
| | Fishing hut ownership | *When I first started going everyone was there for one reason, fishing, and everyone fished. Now … they are just there because it suited their price range and a house and it's an easy trip into town."* [P28] |
| | Local employment | "*If I wanted to restart my guiding career, I don't think I could do it now I certainly wouldn't be able to offer the same level of fishing for them."* [P13] |
| | Community identity | *"People were absolutely passionate about the river and the fishing, there were big fish catches, big numbers of fish being caught so it was, so they were the heydays and it has been going downhill ever since."* [P26] |

### Interview prompts and questions

#### 1. Opening questions

- Can you tell me about your association with this river?
  - Why do you come here?
  - How long have you been coming here?
  - How often do you come here?
  - With whom do you come to the river?
  - What makes this place special?
- Where do you fish along the river?
- What do you enjoy about fishing this river?

#### 2. Location-specific questions and mapping

- Where do you fish along the river?
  - Participatory mapping step: visualise these reaches on the Rakaia River map
- Please estimate the time frames over which you have first-hand experience of the river environment at each location.
- Do you have any historic photos, new articles, recollections or other records (visual or written) that you would be happy to share and reflect the river and fishery over time at these locations?

##### For each location

*Change in the environment/ecology*

- How has the fishing changed over the years ?
- How has the river changed over the years (if at all)?
  - What have you noticed about the water / flows / riverbed / surrounding land-use / vegetation patterns / fish / wildlife / people / weather or climate / seasonality…?
  - Are there any other indicators of change we haven't covered above?
- Are any of these changes associated with benefits, or alternatively concerns, from your perspective? If yes in what way(s)?

*Change in community/people*

- How would you describe the present-day fishing community at this location? How has this community changed in the time you have been associated with this location?
- How do people feel about the state of the river and the fishery at this location (and how has this changed over the years)?
- Who else uses this place?
- What changes have you observed in river user groups over the years that you have been visiting this location?

#### 3. Closing questions on river health and management

- Do you believe the river is in good health? Why/why not? How can you tell?
- Who do you think is responsible, or able to, improve the health of this river?
- What are the most important management actions that would improve the health of the river in your view?
- How would you describe your hopes for the future of this river and fishery?
